# *In vivo* regulation of an endogenously-tagged protein by a light-regulated kinase

**DOI:** 10.1101/2024.11.27.625702

**Authors:** Mengjing Bao, Katarzyna Lepeta, Gustavo Aguilar, Sophie Schnider, Markus Affolter, Maria Alessandra Vigano

## Abstract

Post-translational modifications (PTMs) are indispensable modulators of protein activity. Most cellular behaviours, from cell division to cytoskeletal organization, are controlled by PTMs, their miss-regulation being associated with a plethora of human diseases. Traditionally, the role of PTMs has been studied employing biochemical techniques. However, these approaches fall short when studying PTM dynamics *in vivo*. In recent years, functionalized protein binders have allowed the post-translational modification of endogenous proteins by bringing an enzymatic domain in close proximity to the protein they recognize. To date, most of these methods lack the temporal control necessary to understand the complex effects triggered by PTMs. In this study, we have developed a method to phosphorylate endogenous Myosin in a light-inducible manner. The method relies both on nanobody-targeting and light-inducible activation in order to achieve both tight specificity and temporal control. We demonstrate that this technology is able to disrupt cytoskeletal dynamics during *Drosophila* embryonic development. Together, our results highlight the potential of combining optogenetics and protein binders for the study of the proteome in multicellular systems.

## Introduction

Much progress has been made in the past decades in understanding the molecular and cellular basis of multicellular development (summarized in (Liberali and Schier, 2024)). During this time, much of our understanding of developing systems has relied on the manipulation of development at the genetic level (Lewis, 1978; Nusslein-Volhard and Wieschaus, 1980). This approach has been further boosted by the development of efficient genome engineering technologies (Jinek et al., 2012) and the use of RNA interference (Fire et al., 1998). Despite the dominance of these approaches, the direct manipulation of proteins has attracted increasing attention in recent years. Proteins are the major actors controlling cell behaviour, engaging in complex and dynamic interaction networks. Various methods have been described which allow for a direct control of protein function, including techniques such as anchor-away (Haruki et al., 2008), knock-sideways (Robinson et al., 2010) and the insertion of TEV cleavage sites in a protein of interest, accompanied by time- and space-controlled expression of TEV protease (Pauli et al., 2008). More recently, a large number of optogenetic tools have been developed, allowing protein manipulation in time and space by using light (reviewed in(Emiliani et al., 2022)).

An additional way to directly control protein function has been paved by the isolation and functionalization of protein binders. This type of proteins, to which antibodies and nanobodies belong, are able to recognize with high specificity and affinity a single target protein. Over the past years, many small and genetically-encodable protein binder platforms have been described (Muyldermans, 2021). Protein binders can also be functionalized by fusing them to various effector domains, such as localization scaffolds or enzymes, which will then predictably affect the target protein. The development of protein binders recognizing both endogenous proteins and peptide tags, together with the increasing number of functionalising domains, is generating a powerful toolbox for the direct study of protein function (reviewed in (Schnider et al., 2024),(Cheloha et al., 2020),(Beghein and Gettemans, 2017), (Schumacher et al., 2018),(Frecot et al., 2023),(Ingram et al., 2018).

Cell behaviour depends on sophisticated networks of protein-protein interactions (Low et al., 2021) These interactions are, in turn, dynamically modulated by post-translational modifications (PTMs). Among them, the reversible protein phosphorylation by kinases and phosphatases constitutes an crucial regulatory mechanism that controls multiple cellular processes, such as cell cycle, growth or signaling (Manning et al., 2002). (Cohen, 2002; Hunter, 2000) Aiming to direct protein phosphorylation *in vivo*, we have previously engineered kinases (Synthetic Kinases) that permit tissue-specific phosphorylation of GFP and mCherry fusion proteins (Lepeta et al., 2022). The first synthetic kinase consisted of a fusion of an activated Rho-associated protein kinase (ROCK) to a GFP-binding nanobody. This protein fusion robustly phosphorylates GFP-fused Spaghetti Squash (Sqh), the regulatory light chain of non-muscular Myosin II, when co-expressed in the same cells. Expression of this synthetic kinase via the UAS-Gal4 system allows for the phosphorylation of GFP-tagged Sqh in different tissues, resulting in actomyosin contraction in defined tissues in the developing embryo (Lepeta et al., 2022) .

We reasoned that the utility of synthetic kinases could be further expanded if their activity were to be controlled more acutely, both in time and space. In the past few years, a number of different optogenetic systems that enable temporal and spatial control over biological systems have been developed (reviewed in (Emiliani et al., 2022)). Most interestingly, Leonhard and colleagues recently combined protein-binder-directed targeting of proteins and optogenetic control via light inducible dimerization, in order to trigger degradation in culture cells and in *C elegans* (Deng et al., 2020). Inspired by this study, we used the light dimerization system CIBN/CRY2 to further boost our synthetic ROCK kinase, engineering a light-inducible ROCK which can phosphorylate GFP-fused proteins in a light-dependent fashion. We tested the inducibility and reversibility of the system in developing Drosophila embryos.

## RESULTS

### Design of a light-inducible Rho kinase

With the aim to improve the spatial and temporal control of the phosphorylation triggered by N-Rok:vhhGFP4 (Lepeta et al., 2022), we adapted the optogenetic LiPD modules (Deng et al., 2020), composed by the CIBN/CRY2 light-sensitive dimerization domains. The resulting system, hereof named OptoKinase, is composed by two components: 1) the anti-GFP nanobody VHH4 (also referred as GBP4) (Saerens et al., 2005) fused to the CRY2PHR binding partner 1 (Kennedy et al., 2010), and 2) the activated N-terminal kinase region of Drosophila ROCK fused to the CIBN binding partner 2 (Kennedy et al., 2010) (Figure 1). Upon blue light illumination, the dimerization of CIBN/CRY2 would bring the GFP fusion protein and the synthetic kinase in close proximity, eventually resulting in phosphorylation of the target GFP fusion protein. We generated two Drosophila lines carrying either *UAS-VHH:CRY2* (VhhCRY2) or *UAS-Rok:CIBN* (CIBNRok), controllable by the Gal4/*UAS* system (Brand and Perrimon, 1993).

**Figure. 1.**
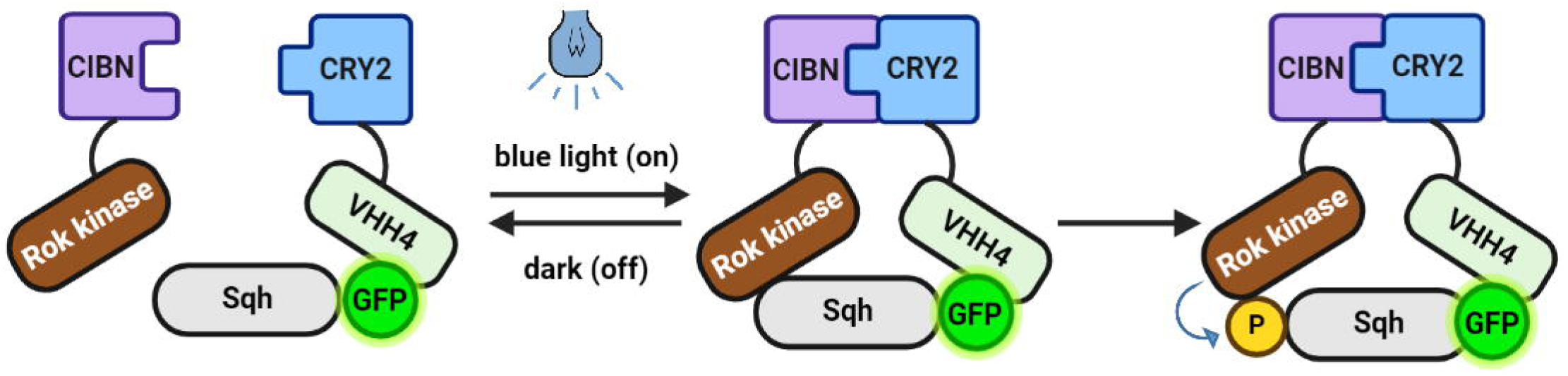
**Schematic illustration of the optogenetic kinases system.**This system is composed of Cryptochrome 2 (CRY2) photoreceptor linked to VHH4 (the anti-GFP nanobody) and the activated Rok Kinase domain fused to the binding partner CIBN (the N-terminal part of CIB1). When induced by blue light, CRY2 dimerizes with CIBN, effectively bringing the constitutively active Rok kinase in close proximity to Sqh::EYFP-HA (here indicated as Sqh GFP fusion). The proximity of the Rok kinase to Sqh::EYFP-HA allows for efficient phosphorylation (P) of Sqh.

In order to test the activity of the OptoKinase, we expressed the *UAS*-driven transgenes in the posterior compartment of each body segment of the Drosophila embryo using the *engrailed-Gal4* (*en-Gal4*) driver. As a target we employed Sqh, endogenously tagged with both HA and EYFP (which can be recognized by the anti-GFP nanobody (Rothbauer et al., 2006)). Using similar *sqh* transgenes as in this setup, we have previously shown that the synthetic Kinase is able to trigger cell contraction and interfere with normal embryonic development (Lepeta et al., 2022).

We first tested whether expression of the minimal catalytic domain of ROCK fused to CIBN would, by itself, result in phosphorylation of Sqh:EYFP-HA. Only minimal changes were observed in Sqh localization pattern or phosphorylation levels, as revealed by anti- phosphoSqh immunostaining, when compared to control (Figure 2B, panels of the first row). In some segments, however, a significant increase in phosphorylation was evident at the leading edge. (yellow arrows) This is in accordance with our previous results (Lepeta et al., 2022), and further supports the dependency on the physical proximity mediated by the nanobody for the minimal kinase to act. Similar results were obtained when the flies were raised in normal cycles of day and night or in total darkness (Fig. 2B first and fourth rows panels).

**Figure. 2.**
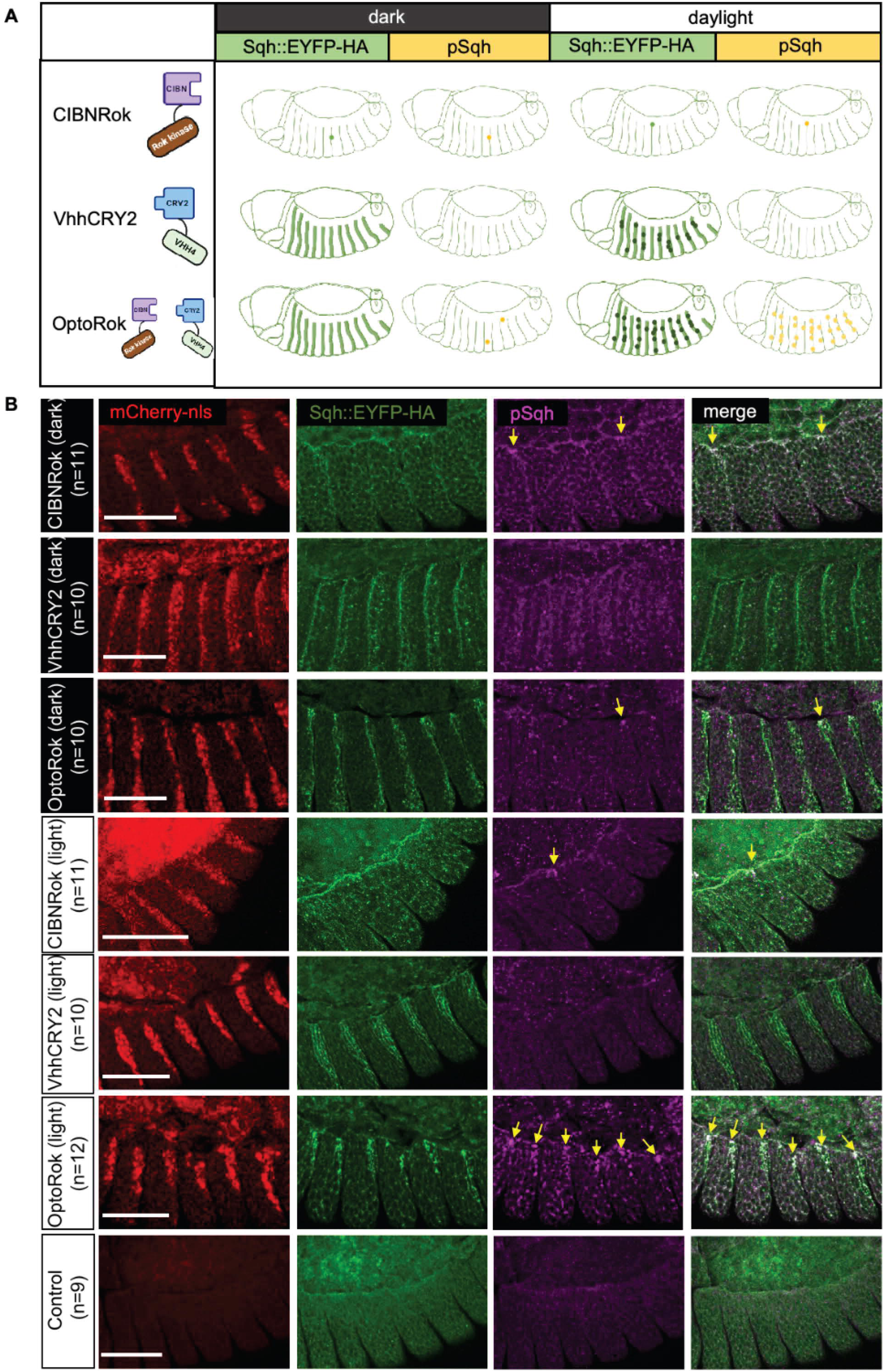
Light-induced OptoRok efficiently phosphorylates Sqh::EYFP-HA in vivo in a tissue-specific manner. A) Schematic illustration of the OptoRok system and expected results on Sqh::EYFP-HA. The UAS constructs are depicted on the left, embryos expressing these constructs with the *en-Gal4* driver are illustrated in the corresponding rows both in the dark or in light conditions. The green stripes represent the fluorescence enhancement of the Sqh::EYFP-HA signal due to the binding of the anti-GFP nanobody (Kirchhofer et al., 2010) and therefore highlighted with VhhCry2 and OptoRok (second and third rows). The dark spots overlapping the green stripes represent an additional “clustering” effect due to both Cry2-CIBN interaction and Cry2-Cry2 dimerization(Bugaj et al., 2013) (Duan et al., 2017), effects which are induced only with blue light exposure. The yellow spots are the foci of Sqh::EYFP-HA (hyper)phosphorylation, visible in the light condition and with OptoRok expression (third row).Random sites of hyperphosphorylation can be observed with the (over)expression of the CIBNRok construct (first row) or in the dark with OptoRok (third row). B) Confocal images with lateral views of fixed *Drosophila* embryos at stage 13–14 (dorsal closure) showing the engrailed expression domain (red nuclei of the posterior part of each epidermal segment), Sqh::EYFP-HA protein (green) and anti-phospho-Sqh immunostaining (magenta). Expressed constructs are indicated on the left of each row, together with the number of considered embryos (n). For the first three rows, flies were raised in total dark condition, while for the last four rows, flies were kept in normal daylight cycles. Yellow arrows point to pSqh foci and the co-localization of pSqh and Sqh::EYFP-HA foci. The embryo genotypes were the following: control: *sqh*^EYFP-HA^/(+). CIBNRok: *sqh*^EYFP-HA^/(+); *en*-*Gal4 UAS*-*mCherry:NLS*/+; *UAS*-Rok-CIBN/+. VhhCRY2: *sqh*^EYFP-HA^/(+); *en*-*Gal4 UAS-mCherry:NLS*/+; *UAS*-VhhCry2/+. OptoRok: *sqh*^EYFP-HA^/(+); *en*-*Gal4 UAS*-*mCherry*:NLS/+; *UAS*- *Rok-CIBN*, *UAS*-*VhhCry2*/+. Scale bar: 50 μm.

Expression of the Vhh4:CRY2 module alone resulted in increased fluorescence of EYFP, which was expected from previous *in vitro* (Kirchhofer et al., 2010) and *in vivo* (Harmansa et al., 2017) studies. The fluorescence enhancement was evident both in normal daylight cycles as well as in darkness (Fig 2B, second and fifth rows panels). In these conditions, Sqh:EYFP-HA was still distributed in the cellular cortex, as is expected for this protein. No evident changes of Sqh phosphorylation were observed.

Finally, Vhh4:Cry2 and CIBN:Rok were co-expressed in the presence of Sqh::EYFP-HA. When these embryos were raised in normal light-darkness cycles, massive hyperphosphorylated clusters were formed in the affected segments, most evident in those cells closer to the leading edge during dorsal-closure (Fig. 2B, sixth row panels). These clusters are highly reminiscent of the ones obtained by the Synthetic Kinase (Lepeta et al., 2022). In contrast, minimal clustering and phosphorylation was observed when the animals were raised in darkness (Fig. 2B, third row panels). A schematic representation of these findings is depicted in Fig. 2A. These results demonstrate that the minimal kinase domain can phosphorylate the endogenous Sqh upon dual recruitment by both light and the interaction of the nanobody and EYFP.

### Light-induced Sqh hyperphosphorylation disrupts dorsal closure

We next investigated the effects of the light-inducible Sqh phosphorylation during the morphogenetic process of dorsal closure by imaging the above-mentioned genotypes and subjecting the embryos to different protocols of illumination. First, we collected and maintained all embryos in ambient light, which we referred as daylight, and then transferred them to the microscope room and started imaging as described.

Under these conditions, we expected that dimerization of the two components, if present, would be induced at all times. Control *sqh*^EYFP-HA^ embryos (Figure 3A, first row), expressing none of the two optogenetic components, closed the dorsal opening in about two hours, and exhibited an uniform actomyosin cable in the epidermal leading edge cells (revealed by accumulation of Sqh::EYFP-HA) and a regular pairing of epidermal segments (see also movie S1). This is similar to what was observed in control, wild-type embryos (see(Lepeta et al., 2022)).

**Figure 3.**
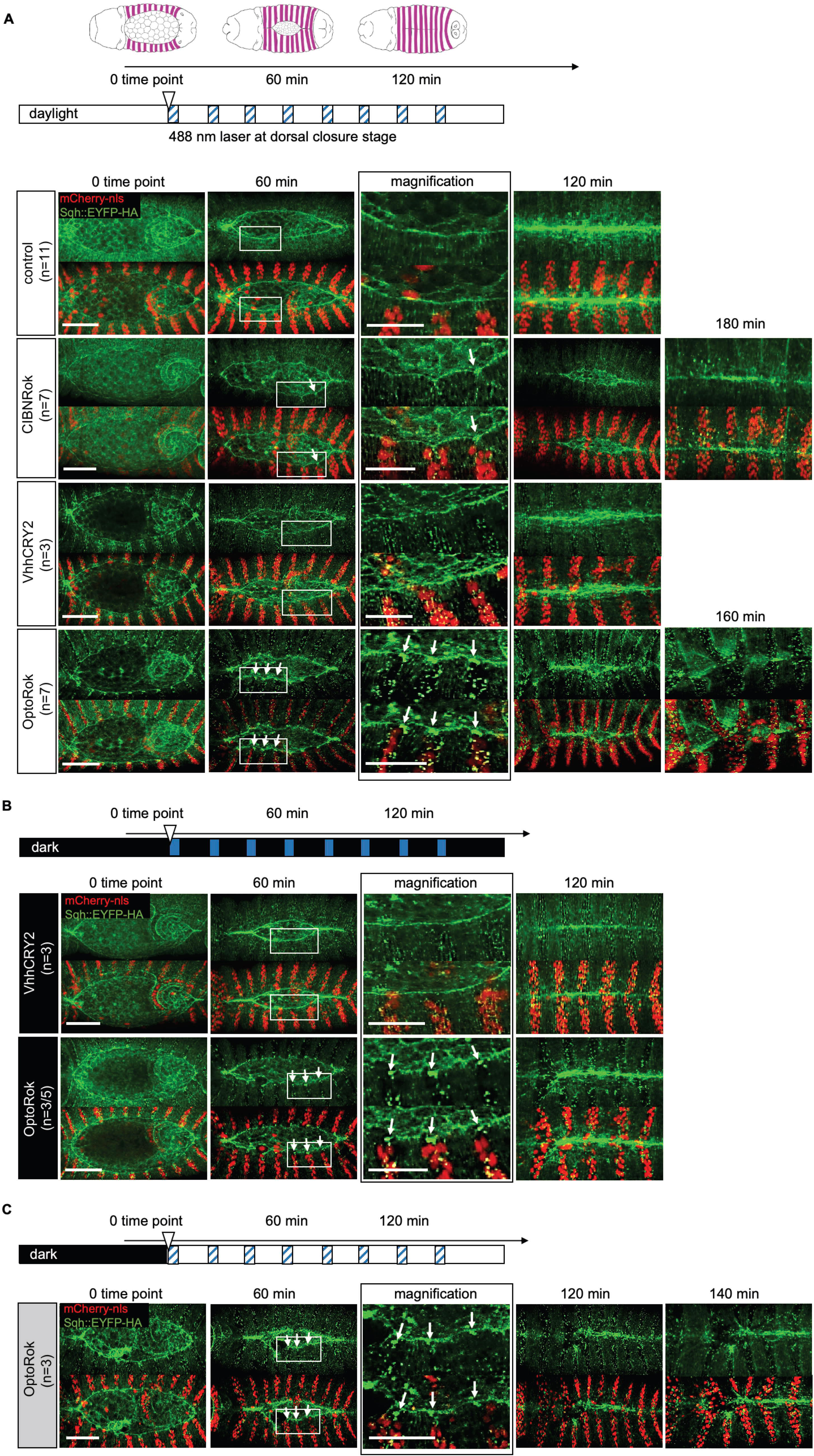
Light-induced OptoRok kinase modulates mechanical properties of cells through the phosphorylation of Sqh::EYFP-HA and myosin II activity. A) Schematic illustration of the dorsal closure process in the developing fly embryo. Dorsal closure was used as a model to assess myosin II activation using OptoRok kinase. The entire process, including fly crossing, embryo collection, dechorionation, and imaging, was conducted under ambient light. The clear horizontal bar represents the imaging process, with the starting 0 time point indicated by a clear triangle and the acquisition times of about 3 min with the 488nm laser for the green channel indicated by the blue striped boxes. These extra “blue” light stimulations were repeated every 20 minutes over the whole process of dorsal closure. All panels show stills from live-imaging with dorsal views of the developing embryos at stages 13/14/15 (dorsal closure) visualized by Sqh::EYFP-HA (in green) and expressing CIBNRok, VhhCRY2 and OptoRok (constructs indicated on the left, together with the number of considered embryos (n) in the engrailed domain (red nuclei with mCherry-nls). The magnification panels (third column) are from the 60 min time point. The white arrows are pointing to the Sqh::EYFP-HA foci and actomyosin cable invaginations at the 60-minutes time point and the magnification in both CIBNRok and OptoRok panels. CIBNRok and OptoRok showed longer time for dorsal closure (up to 180 minutes, 5th column) than control. The genotypes are the same as in Fig. 2B, except the control: *sqh*^EYFP-HA^/(+); *en*-*Gal4 UAS-mCherry:NLS*/+ B) Fly crossing, embryo collection, and dechorionation were performed under red light, equivalent to darkness. As indicated by the black horizontal bar, the only light stimulation was during the 3 min acquisition time with the 488 laser (blue boxes) every 20 minutes, as the low ambient light conditions of the microscope room were reduced to minimum. Expressed constructs, genotypes and panels arrangements are the same as in A). C) Same as in B) but maintaining the low ambient condition of the microscope room during acquisition. The n number of OptoRok embryos was given as the proportion of embryos showing a slightly aberrant closure pattern at the 120-minute time point relative to the total number of embryos imaged. The remaining embryos exhibited a normal closure pattern. Scale bar: 50 μm; for the magnification panels: 25 μm.

In embryos expressing CIBNRok (second row, Fig. 3A), a moderate effect on the dorsal closure process was observed. This was manifested in local invaginations of the cable, occasional formation of Sqh::EYFP-HA foci (white arrows), and in a delay in closure of up to 3-4 hours. These findings were consistent with the increased Sqh::EYFP-HA phosphorylation seen in some segments upon CIBNRok expression (see Fig. 2B).

VhhCRY2 expression caused a light-dependent enhancement of Sqh::EYFP-HA fluorescence within the posterior compartments, a phenomenon that we had already observed in the fixed embryos (Fig. 3A, third row). We also observed slight distortions of the cable structure, maybe due to partial Sqh::EYFP-HA clustering induced by Cry2 oligomerization (Bugaj et al., 2013), but embryos completed dorsal closure with regular timing and exhibited an almost regular epidermal patterning.

Expression of OptoRok in this constant illumination protocol resulted in a significant phosphorylation of Sqh::EYFP-HA (shown in Fig.2 ) and formation of prominent Sqh::EYFP- HA foci (Fig. 3A, fourth row). In comparison with RokCIBN alone, the foci were noticeably larger, clearly visible already at earlier stage of closure (Fig. 3A, first column) and present along the entire *enGal4* expression domain, and not only in the cells surrounding the dorsal gap. The presence of Sqh::EYFP-HA foci lead to local invaginations of the actomyosin cable, delayed closure, and - in sharp_contrast to the control condition - to mispairing of the epidermal segments resulting in disrupted or, in some cases, failed closure (Fig. 3A, fourth row; see also Movie S2). These effects mimicked those we observed with N-Rok::vhh4GFP4 (Lepeta et al., 2022), although to a somewhat lesser degree, and were similar to what we observed with N-Rok::dGBP1 fusion that harbors a destabilized GFP-binding nanobody (dGBP1(Tang et al., 2016).

We next tested whether different illumination protocols could modulate the severity of the observed effects. When the embryos were kept in the dark all the time, including reducing at minimum the ambient light in the microscope room and only stimulated during the acquisition phases (about 3 min with 488 laser every 20 min) (Figure 3B), we observed formation of some Sqh::EYFP-HA foci (white arrows) and a distortion of the cable structure, indicating activation of OptoRok (second row, Figure 3B). This activation occurred rapidly, as we observed the foci already at time point 0 to 3 minutes, but generally the embryos closed in the normal time and presented only minor closure defects (Movie S3).

Conversely, when the embryos kept in the dark were subjected to the low ambient light during the entire imaging process (in addition to the extra 488 stimulation during the acquisition times (Fig. 3C and Movie S4), we observed the same phenotype and delay in dorsal closure as for those exposed to constant daylight, i.e. dorsal closure defects, including the formation of Sqh::EYFP-HA foci, local invaginations of the cable structure at these foci, delayed closure, and deformation of the epidermis. These experiments indicate that different illumination protocols can indeed tune the functionality of the OptoRok system, which, in turns, may modulate the *in vivo* mechanical properties of cells by regulating the phosphorylation status of Sqh::EYFP-HA.

### Light-induced hyperphosphorylation of Sqh persists in the absence of light

A useful property of many optogenetic systems, besides their spatial and temporal inducibility, is their reversibility(Bugaj et al., 2013; Kennedy et al., 2010; Levskaya et al., 2009; Wu et al., 2009). In order to test if the hyperphosphorylation of Sqh persisted in the absence of light, 18 hours old embryos kept in normal light-darkness cycles were either maintained in ambient light or placed in darkness. Embryos were subsequently incubated for one- or two-hours prior fixation (Figure 4). In all these conditions, hyperphosphorylation of Sqh, measured by anti-p-Sqh antibody staining, was consistently high and concentrated in clusters (when compared to the cells where the OptoRok was not expressed (Fig. 4, yellow arrows in the magnification panels)). This result indicates that Sqh hyperphosphorylation is not immediately reversible under these conditions.

**Figure 4.**
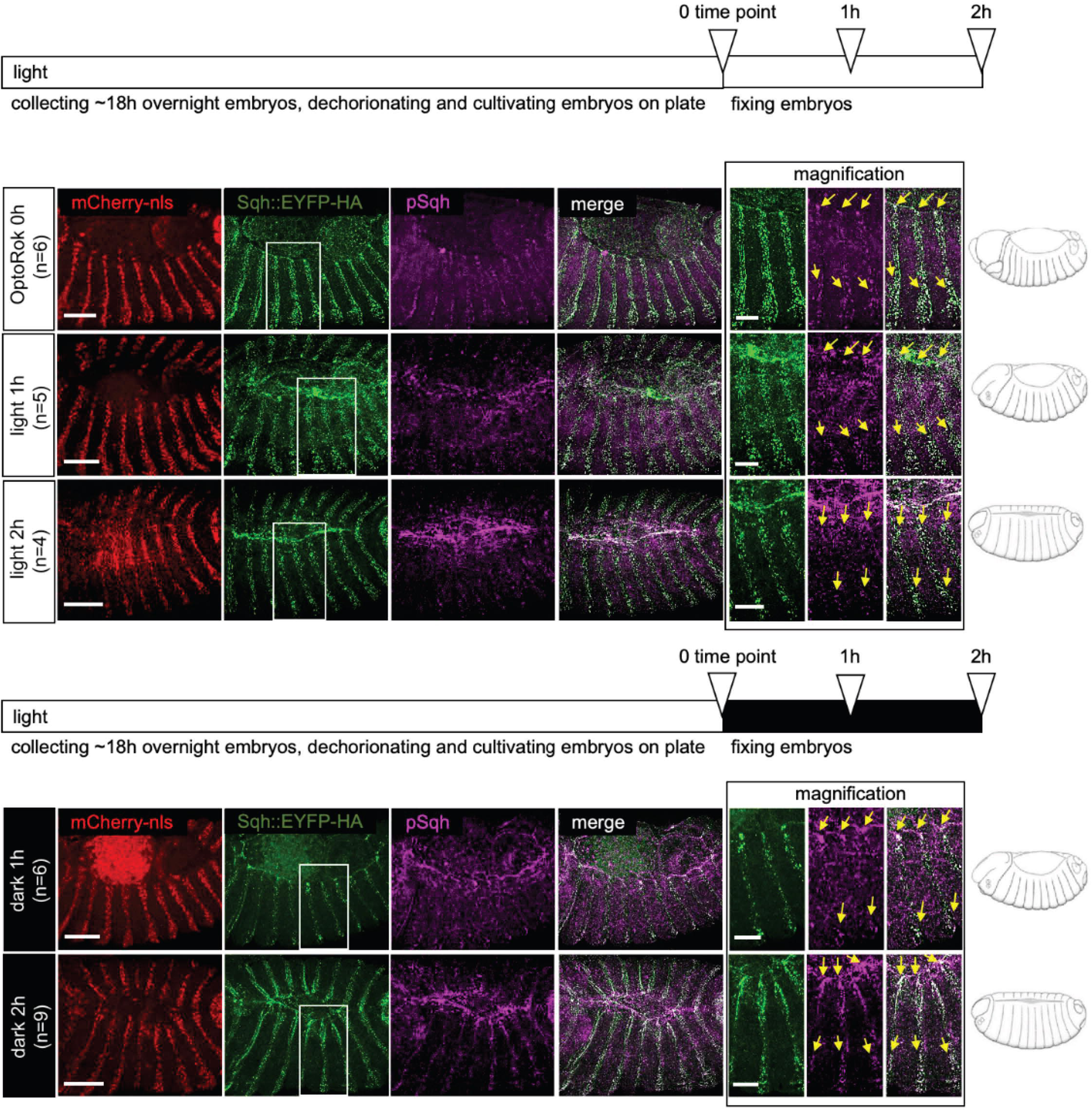
The light-induced phosphorylation of Sqh::GFP by OptoRok remains stable in the dark during the dorsal closure period. Fly crossing, embryo collection, and dechorionation were conducted under standard light-darkness cycles. As indicated in the horizontal bars, at 0 time point , ∼ 18hours-old embryos were either collected or transferred to fresh plates to continue development in plain ambient daylight (first three rows) or in the dark (fourth and fifth rows). After 1 or 2 hours (open triangles on the horizontal bar) embryos were collected, fixed and stained with anti-phospho-Sqh antibody. Embryos exhibiting morphological patterns characteristic of the relevant stage (schematically depicted on the right of each row) were selected. Confocal images of embryos with this genotype: *sqh*^EYFP-HA^ /(+); *enGal4 UAS-*mCherry:NLS/+; *UAS-Rok-CIBN*, *UAS-VhhCry2*/+ are shown. The colour scheme is the same as in Fig. 2. A magnification of about 3 epidermal segments , including the cable structure, is at the right of each row.. Yellow arrows point to pSqh. Scale bar, 50 μm, for the magnification panels, 25 μm.. Note that at the 2-hour time point, the dorsal closure of OptoRok is not yet complete, as shown in Fig. 3. We, therefore, selected nearly closed embryos as the 2-hour time point.

## Discussion

Controlling the post-transcriptional modification of proteins *in vivo* is indispensable to dissect the functional role of these PTMs. Along those lines, we have previously introduced engineered kinases which allow the directed phosphorylation of GFP-tagged targets (Lepeta et al., 2022). In order to improve their precision, we thought of using optogenetic dimers, thus adding light inducibility and possibly reversibility to their enzymatic reaction. Based on elegant previous studies published by the group of Heinrich Leonhardt (Deng et al., 2020), we opted for the use of the CIBN/CRY2 system and fused the activated Rho kinase to CIBN and the GFP-binding nanobody VHH4 to CRY2. Upon photoactivation, the activated kinase is recruited to the GFP-fused target protein (Sqh::EYFP-HA in our case) via the kinase-associated GFP nanobody, which eventually results in Sqh::EYFP-HA phosphorylation and actomyosin contraction.

Although the system allows for optical induction and the subsequent phosphorylation of Sqh::EYFP-HA, it turns out that when the activated kinase fused to the GFP nanobody (such as initially described in (Lepeta et al., 2022)) is split into two parts by the photodimerization domains, the kinase is somewhat less active. As described previously, activated Rho kinase phosphorylated Sqh::EYFP-HA very effectively as visualized by phospho-Sqh antibody staining and by clustering of Sqh::EYFP-HA in cells expressing the kinase (Lepeta et al., 2022). Although we do see pSqh accumulation (Figs. 2B and 4), we see weaker effects on dorsal closure with OptoRok than with N-Rok::vhh4GFP4. This might indeed be due to the splitting of the nanobody-RhoK construct into two portions, the CRY2 and the CIBN parts. This larger molecular complex formed upon light on the GFP target protein might put the kinase moiety at a more distant position and render it somewhat less active. Nevertheless, these “weaker” effects were more similar to those observed with to Rok::dGBP1, the optimized variant with the destabilized nanobody (Lepeta et al., 2022) (see in the next paragraph). It is also worth noting that some phenotypic differences may be also due to the sex of the embryos, since female embryos have one copy of untagged *sqh*..

The fusion of the light-inducible dimerization domains to Rok and the VHH4 nanobody, respectively, results in hybrid proteins which also have activities by themselves. We observed that the expression of only the VhhCRY2 part induces enhancement of EYFP fluorescence; it has previously been shown that binding of the GFP nanobody to GFP does increase fluorescence (Harmansa et al., 2017; Kirchhofer et al., 2010; Lepeta et al., 2022), so the enhancement we see is unlikely to reflect an increase in the amount of the GFP fusion protein but rather an increase in fluorescence, and this effect is light-independent (Figure2B and 3). This signal enhancement might also reflect a partial clustering effect due to the Cry2 module light-induced oligomerization (Bugaj et al., 2013). Interestingly, this particular property was exploited in a recent paper for developing an optogenetic trapping tool (Xu et al., 2024). The expression of the CIBNRok fusion leads to a slight increase in pSqh levels (Figure2B), and a slight increase in the time it takes to close the dorsal opening of the developing embryo (Figure3A). Since the Rok construct used in this fusion protein carries activating mutations (see (Lepeta et al., 2022)), occasional close proximity of Sqh and CIBNRok may lead to phosphorylation of the former. To circumvent this problem upon the expression of the direct fusion construct VHH4Rok, we replaced VHH4 with a conditionally stable nanobody; only upon binding to GFP the fusion protein was stabilized, reducing the background activation levels in the absence of the target (Lepeta et al., 2022). Such a strategy is not possible with the CIBN/CRY2 system, since both proteins must be stably expressed in the absence of light in order for them to dimerize upon light administration.

We also tested the reversibility of the light-induced phosphorylation and found that the system is not reversible on a short time scale. The half-dissociation time of the CIBN/CRY2 complex is approximately 5-8 min (Deng et al., 2020). Assuming similar kinetics in the Drosophila early embryo, after one hour in the dark, most of the OptoRok should be separated into its two components and the kinase should thus not be recruited to the target GFP fusion protein anymore (see Fig 1); however, we still see similar P-Sqh levels after two hours in the dark to the levels seen in activating (light) conditions (Figure4). We think that this is most likely due to insufficient amount/activity of phosphatases which would remove the excess phosphate added by the ectopic kinase expression. It is thus possible that reversibility of actomyosin contractility can only be obtained by simultaneous expression of the corresponding phosphatases.

When following the effects of photoactivation via live imaging, it would be preferable to monitor the developmental consequences via a fluorescent protein marker which can be visualized in spectral light condition which do not activate dimerization, e.g. a red fluorescent protein, or proteins visible in the red, far-red or infrared spectrum. In our scenario, we could monitor the effects on dorsal closure in a genetic background in which one endogenous copy of Sqh is tagged with mCherry. We have recently generated such flies and are planning to use them in similar experiments as those described here. It would also be possible to use other tagged proteins as markers, such as RFP-Moesin, RFP-actin or RFP-E-cadherin. These different settings would also allow for a clean light inducibility, monitoring in real time the appearance of the phenomenon (in our case Sqh clustering) after illumination, from a zero time point where no clusters are present. Another strategy could be a very precise optimization of imaging parameters to image EYFP without activating the optogenetic system. EYFP has an excitation peak around 513 nm. By avoiding shorter wavelengths and using a narrow-band filter that specifically excites EYFP, the likelihood of exciting the blue light-sensitive optogenetic system might be strongly reduced (Kennedy et al., 2010) .

As a general comment, an enzymatic activity such as phosphorylation can benefit from an optogenetic “module”, because one can control its induction in time and space very precisely. However, if other enzymes are needed to remove the results of its activity (such as the phosphatase in the case of kinase activation), the reversibility might not be easily applicable.

In summary, more work is needed to generate a more powerful optically regulated actomyosin toolbox using protein binder tools.

## Aknowledgements

We would like to thank the Biozentrum Imaging core facility for their great help and support, Heinrich Leonhardt (Ludwig-Maximilians-Universität München, Munich, Germany) for the piggyBac-LIPD -GBP-IR plasmid and Robert Ward (Case Western Reserve University, Cleveland, OH) for the anti Psqh antibody.

We are grateful to Karin Mauro, Bernadette Bruno, Maria del Consuelo Zuluaga Gomez and Gina Evora for constant and reliable supply of fly food and to all members of Affolter lab for the helpful discussions.

The work in the laboratory of M. Affolter was supported by grants from the Swiss National Science Foundation (310030_192659/1) and by funds from the Kantons Basel-Stadt and Basel-Land.

## MATERIALS and METHODS

### Plasmid construction

The optogenetic plasmids were generated by specific PCR amplification and standard restriction cloning using the pUASTattB vector (Bischof et al., 2007). Briefly, the pUASTattBVhhGFP4-Cry2 was constructed by inserting a Xba1/BstE2 Cry2-myc fragment from piggyBac-LIPD -GBP-IR (Deng et al., 2020) into pUASTattBVhh4 (Caussinus et al., 2011). For pUASTattBRhokinase-CIBN, the CIBN-HA fragment was PCR amplified from the same piggyBac-LIPD -GBP-IR and inserted Eag1/Xba1 into pUASTattB_N-Rok::dGBP1 (Lepeta et al., 2022). The constructs were all verified by sequencing.

### Fly injection

Fly embryos were injected as follows: 30 min old eggs were dechorionated in 3.5% bleach solution, aligned using a stereo microscope and adhered on a glass slide. To avoid desiccation, embryos were covered with Voltalef H10S oil. PBS diluted plasmids were then injected in the posterior pole using a glass needle with the help of a pressure pump and a micromanipulator. Each plasmid, at a concentration of 100ng/μl, was injected in embryos containing *nos-phiC31* and one of the following third chromosome attP landing sites: ZH- 86Fb for pUASTattBVhhGFP4-Cry2 and {3xP3-RFP}ZH-64A for pUASTattBRhokinase-CIBN.

### Fly genetics and husbandry conditions

All flies used in this study were grown in regular polenta/yeast vials and kept at 25° C unless specified and subjected to regular day/night cycles of 12 hours. Embryos collection, manipulation and imaging were performed in ambient light during the 12h daylight period. For the dark condition, flies were kept in a lightproof box from the cross setting on and embryos were collected and mounted under only a red light source.

### Fly stocks generated and employed in this study

w *sqh*^EYFP-HA^ generated in this lab by CRISPR/CAS methodology and described in detailed elsewhere (Schnider et al, in preparation) (Aguilar et al., 2024) w *sqh*^EYFP-HA^; *enGal4 UAS-mCherryNLS* (Lepeta et al., 2022) w; M{*UAS*-Vhh-Cry2, *w*^[+]^}zh-86Fb w; M{*UAS*-Rok-CIBN, *w*^[+]^}zh-64A w; M{*UAS*-Rok-CIBN, *w*^[+]^}zh-64A M{*UAS*-Vhh-Cry2, *w*^[+]^}zh-86Fb

### Immunostaining of Drosophila embryos

Embryos were collected on grape juice agar plates supplemented with yeast paste after overnight egg laying at 25°C and processed for immunofluorescence as in (Lepeta et al., 2022). Briefly, embryos were collected, dechorionated in 3.5% bleach (sodium hypochlorite, stock solution 13% w/v technical grade; AppliChem GmbH), washed thoroughly with H2O, and fixed in 50:50 heptane: 4% paraformaldehyde solution for ∼20 min with vigorous shaking. Embryos were blocked in PBTN (PBS + 0.3% Triton X-100 + 2% normal goat serum) for 1 hour, and incubated overnight with primary guinea pig anti-Sqh1P antibody (gift from R. Ward; (Zhang and Ward, 2011)) at 1:400 concentration at 4°C and secondary antibodies Alexa Fluor 647 (1:500; Thermo Fisher Scientific) for 2 hours at room temperature. Embryos were mounted on a microscope slide in Vectashield mounting medium (H-1000; Vector Laboratories) and covered with a 22-mm^2^ coverslip.

### Embryo mounting for live time-lapse imaging

Embryos were dechorionated in 30–50% bleach (see above) and extensively rinsed with water. Embryos at the desired stages were manually selected on a grape juice agar plate under a dissecting microscope and mounted on a glass-bottom dish (MatTek, 35 mm dish, no. 1.5 coverslip, uncoated, P35G-1.5-10-C). After adding 1x PBS, the embryos were gently rolled into the desired positions using a cut gel loading pipette tip. Properly dechorionated embryos adhered to the glass bottom, with the part to be imaged facing down for the inverted microscope.

### Confocal and time-lapse imaging

Images of fixed embryos were acquired with 1,024 × 1,024 frame size with a PLAN APO 40×/1.2NA objective (LD LCI PLAN APO, Imm Corr DIC M27, water immersion) on Point Scanning Confocal Zeiss LSM880. A series of z-stacks was acquired for each embryo at 0.2–0.6 μm step size and a 488, 561, and 633 nm laser line. Time-lapse sequences of dorsal closure were imaged under Point Scanning Confocal Zeiss LSM880 inverted microscope with a PLAN APO 40×/1.2NA objective. A series of z-stacks were acquired for each embryo at 0.3– 1 μm steps using a 488 and 561 nm laser line. Imaging was carried out at 20 min intervals. Z-stack maximum projections were assembled in Fiji (Schindelin et al., 2012) or OMERO (Allan et al., 2012) Figures were prepared using Adobe Illustrator.

### Illumination protocols

We refer to Daylight the ambient light of the laboratory during the day, which have both natural light from windows and artificial standard white light. The microscope room has a very low ambient light which could be further reduced by covering the instruments with a black cloth.

During the imaging sessions, samples receive 3 minutes pulses with the 488 nm laser for the acquisition of the green channel, repeated every 20 minutes over the whole process of dorsal closure.

---

**Movie S1.** The control embryo shows normal dorsal closure.

**Movie S2.** Expression of OptoRok in embryo under light exposure leads to abnormal dorsal closure.

**Movie S3.** Expression of OptoRok in embryo in the dark shows normal dorsal closure.

**Movie S4.** Expression of OptoRok in embryo kept in the dark followed by light activation during imaging results in abnormal dorsal closure.

The color scheme and the genotype of the embryos shown in the movies are described in Fig.3 legend.

